# EEG Dynamics of Movement Preparation and Error Processing Distinguish Motor Adaptation From De Novo Learning

**DOI:** 10.64898/2025.12.11.693740

**Authors:** Raphael Q. Gastrock, Denise Y. P. Henriques, Bernard Marius ’t Hart

## Abstract

Previous research has distinguished behavioral mechanisms between motor adaptation and de novo learning. However, electroencephalography (EEG) markers dissociating them remain unclear. Here, participants (N = 32, 13 female) performed center-out reaching under a 30° fixed rotation (adaptation), mirror reversal (de novo learning), or random rotation (errors without learning), while we recorded EEG during movement preparation and post-movement feedback. We explored how perturbation type, training phase, and error magnitude shaped neural dynamics in both temporal and frequency domains. During preparation, the readiness potential (RP) showed opposite training-related modulation in the fixed and random rotations, suggesting increased reliance on updated internal models for adaptation. Mirror reversal training showed no RP change, likely reflecting reliance on effortful, explicit learning. Interestingly, we found that Lateralized Readiness Potentials (LRPs) reflected planned movement direction rather than effector-specific preparation. In the frequency domain, beta attenuation for large errors and alpha synchronization for small errors were more pronounced in the fixed rotation than in the mirror reversal, suggesting greater error-sensitivity during adaptation. During post-movement feedback, P3 amplitude decreased from early to late learning in the fixed and random rotations but remained stable in the mirror reversal, highlighting differences in cognitive demands across perturbations. A sustained late positivity emerged following small errors across perturbations, potentially indexing implicit learning. Random rotations also elicited distinct frontal and central beta modulation, likely reflecting disengagement due to task unpredictability. Together, we show that preparatory and feedback-related EEG signatures differ across perturbation types, revealing distinct neural mechanisms underlying motor adaptation and de novo learning.

**Significance statement:** Understanding how the brain supports skill acquisition and adaptation is critical for advancing theories and applications of motor learning. Although behavioral differences between motor adaptation and de novo learning are well established, the neural processes that distinguish them remain unclear. By comparing EEG activity during movement preparation and feedback-error processing across multiple perturbation types, we uncover distinct temporal and frequency domain signatures that uniquely characterize each learning process. Our findings show that adaptation engages neural dynamics consistent with updating internal models and error-processing, whereas de novo learning relies on explicit, cognitively demanding strategies that likely unfold over extended practice. These results provide a clear neural framework for distinguishing motor learning mechanisms.

## Introduction

When people encounter movement errors, they adjust subsequent plans. This error processing is central to de novo learning and motor adaptation: de novo learning establishes new control policies, whereas adaptation modifies existing policies to restore performance (Krakauer et al., 2000; Bastian, 2008; Shadmehr et al., 2010; Telgen et al., 2014; Krakauer et al., 2019). Behavioral differences between these two are well-documented (Bock et al., 2001; Werner & Bock, 2010; Gutierrez-Garralda et al., 2013; Telgen et al., 2014; Wang & Taylor, 2021; Wilterson & Taylor, 2021; Gastrock et al., 2024). Adaptation is also well-studied with EEG (Reuter et al., 2022), yet no work has directly compared EEG dynamics between adaptation and de novo learning. Using the paradigm from Gastrock et al. (2024), where participants trained with both visuomotor rotation and mirror reversal perturbations, we explore EEG markers during movement preparation and feedback that distinguish these motor learning types, offering insight into how the brain prepares actions and processes errors.

Different behavioral measures distinguish adaptation from de novo learning. In adaptation, compensation for visual or mechanical perturbations reflects forward model updates driven by sensory prediction errors - discrepancies between predicted and actual outcomes (Shadmehr & Mussa-Ivaldi, 1994; Blakemore et al., 1998; Krakauer et al., 2000; Bastian, 2008; Ostry et al., 2010; Haith & Krakauer, 2013; Krakauer et al., 2019). These updates produce reach aftereffects when the perturbation is removed, indicating implicit adaptation. In contrast, mirror reversal de novo learning, where feedback is flipped across an axis, yields no aftereffects (Telgen et al., 2014; Wilterson & Taylor, 2021; Gastrock et al., 2024). Instead, individuals switch control policies, likely relying on explicit processes. Mirror reversal learning is more cognitively demanding, showing longer preparation times, greater movement variability, and requiring more practice, but yields strong retention and broad generalization (Gastrock et al., 2024). Such differences suggest distinct underlying neural mechanisms between these perturbations.

Event-related potentials (ERPs) capture temporal-domain neural dynamics. The Readiness Potential (RP), a slow-developing fronto-central negativity preceding self-initiated actions, reflects preparatory activity and processing of expected sensory consequences (Kornhuber & Deecke, 1965; Jo et al., 2014; Reznik et al., 2018; Vercillo et al., 2018; Wen et al., 2018; Pinheiro et al., 2020). Effector-specific preparation appears in the Lateralized Readiness Potential (LRP; Smid et al., 1987; de Jong et al., 1988; Gratton et al., 1988), which shows stronger negativity contralateral to the moving hand, though most RP/LRP research uses button presses rather than reaches. Given reaction-time differences between adaptation and de novo learning (Gastrock et al., 2024), these components may vary across perturbations. Following movement errors, fronto-central ERPs show a Feedback-Related Negativity (FRN; Maurer et al., 2019, 2021; Reuter et al., 2020, 2022), followed by a P3 component that scales with error size and remains elevated with reduced explicit learning (Aziz et al., 2020; Palidis et al., 2019; Reuter et al., 2020), consistent with its role in cognitive and attentional processing (Verleger, 2020). Here, we test how error size and learning modulate RP/LRP, FRN/P3 across perturbations.

Frequency-domain EEG dynamics reflect band-specific synchronization. Adaptation studies focus on beta, alpha, and theta (Reuter et al., 2022). Beta decreases after large errors and recovers as errors reduce (Kilavik et al., 2013; Tan et al., 2014; Ozdenizci et al., 2017). Error-driven beta attenuation over contralateral motor regions reflects implicit adaptation, whereas medial frontal beta relates to explicit, strategic re-aiming (Torrecillos et al., 2015; Jahani et al., 2020). Frontal alpha and theta resynchronize with improved performance, reflecting changes in task engagement and cognitive control respectively (Gentili et al., 2011; Arrighi et al., 2016; Savoie et al., 2018; Desrochers et al., 2020). We compare synchronization patterns in these frequencies across frontal and contralateral central regions during perturbation training.

This study extends distinctions between adaptation and de novo learning by examining temporal and frequency-domain EEG during pre- and post-movement phases. Participants trained with a fixed 30° visuomotor rotation, mirror reversal, and a random rotation where participants encounter unpredictable errors. We hypothesize behavioral and EEG dynamics to distinguish across perturbation types, with error magnitude and learning modulating neural activity across domains.

## Methods

They waited for a go signal (target turned blue) before moving to the cued target location. The cursor remained visible throughout the reach, and they were instructed to hold their terminal position.

### Participants

32 healthy adults (13 female, *M*Age = 21.6, *SD*Age = 3.3) participated in the experiment. All participants reported being right-handed, had normal or corrected-to-normal vision, and had no known neurological disorders. We used the same experimental task as in our previous study (see study 1, Gastrock et al., 2024), so we collected the same sample size in the current study. All participants gave written informed consent prior to participating, and received monetary compensation (CAD$ 20/ hour) or course credit for their participation. All procedures were in accordance with institutional and international guidelines, and were approved by York University’s Human Participants Review Committee.

### Experimental set-up

Details for the apparatus, stimuli, and trial types are similar to Gastrock et al. (2024), with some modifications to technical details to incorporate EEG recordings.

### Apparatus

Participants sat on a height-adjustable chair with a digitizing tablet (Wacom Intuos Pro, 16.8” x 11.2” x 0.3”, resolution resampled to 1024 × 768 pixels at 60 Hz) directly in front of them and a vertically mounted monitor (Dell Computer, 19” Ultrascan P991 CRT) positioned 38 cm ahead of the tablet (Fig. 1A). Using their right hand, they moved a digitizing stylus across the tablet surface within a circular stencil (5.1 cm radius, see OSF: ’t Hart et al., 2024) that constrained stylus movement. Head and arm positions were unconstrained, but we instructed participants to sit comfortably and minimize extraneous movements within a given trial.

**Figure 1.**
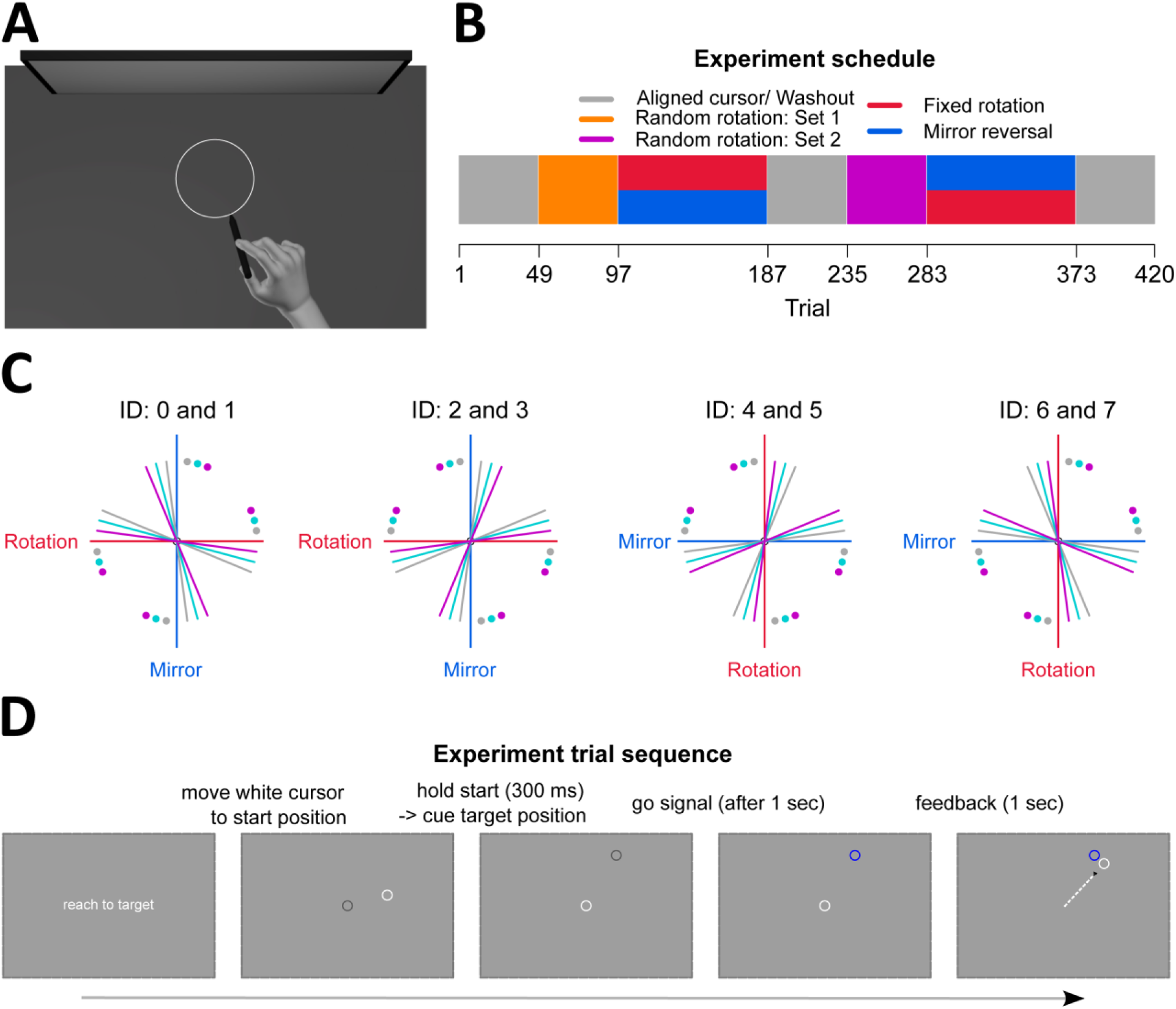
Experimental set-up. (**A**) Participants used a stylus to move across a digitizing tablet, while a monitor displayed stimuli and visual feedback of their hand position. Movements across the digitizing tablet were restricted by a circular stencil (white circle). (**B**) Aligned cursor, or baseline, reaches had matched cursor and hand positions. Participants (N = 32) completed 48 aligned cursor trials, followed by 48 random rotation trials, 90 training trials with either rotation or mirror reversal perturbations, and 48 washout trials. They then completed 48 random rotation trials, 90 training trials with the other perturbation, and 48 washout trials. (**C**) Even numbered participant IDs experienced the rotation before the mirror reversal, and vice-versa for odd IDs. Each perturbation type corresponded to either the vertical or horizontal midline axis and was counterbalanced across participants. Each axis had six target locations, either 7.5°, 15°, or 22.5° away from the axis in the clockwise or counterclockwise direction, and was also counterbalanced across participants. Colored dots indicate targets, and solid lines show the corresponding correct hand movement directions. (**D**) Participants moved the stylus to bring a white cursor to the centre start position.

### Stimuli

A white cursor (1 cm diameter) represented the stylus position on the monitor. Participants made ballistic reaches from the grey start position (1 cm diameter at screen center) toward a blue target (1 cm diameter) until they hit the circular stencil (Fig. 1A, 1D). Targets were radially arranged 4.58 cm from the start position (target center 0.52 cm away from the stencil). Target locations depended on participant ID (Fig. 1C), which determined perturbation order, axis orientation, and perturbation direction, enabling counterbalancing across conditions while avoiding systematic biases in behavior and EEG measures.

Each participant trained with both perturbation types: even IDs experienced the rotation before the mirror task and vice-versa for odd IDs. Perturbations were applied across either the vertical or horizontal midline axis. Each axis and perturbation type had six corresponding training targets (7.5°, 15°, or 22.5° away from the axis, clockwise or counterclockwise direction). This design produced eight counterbalanced conditions (2 perturbation orders X 2 axis orientations X 2 perturbation directions; Fig. 1C). The six targets were presented once in a shuffled order before repeating, ensuring evenly distributed reach directions across locations. Trial types proceeded in a particular order: familiarization, aligned trials, random rotation trials, and perturbed trials (Fig. 1B). The random rotation and perturbed trials were then repeated for the second perturbation type. Each set of perturbation trials is also followed by aligned washout trials. In combination with the randomly rotated trials, this ensures that learning for one perturbation type does not affect the other. Consistent with prior work, we observe aftereffects following rotation, but not mirror reversal training (see R notebook, Gastrock et al., 2025). However, as our focus is on learning during perturbation training, we do not discuss washout results here.

### Trial types

*Familiarization*. Participants kept the hand-cursor at the start position for 300 ms (Fig. 1D, panels 2-3). A grey circle (1 cm diameter) cued the target location and remained for one second as participants continued to hold their position. After this hold period, the target cue turned blue, acting as the go signal to start moving (Fig. 1D, panel 4). Participants moved with continuous cursor feedback until they sliced through the target and hit the stencil. They were then instructed to hold their endpoint position for one second (Fig. 1D, panel 5). To ensure ballistic movements, participants had to reach through the target within 400 - 700 ms of the go cue. Additionally, they had to keep the cursor within 1 cm of the reach endpoint for one second. During familiarization trials only, we required participants to satisfy both criteria for the target to disappear and the start position to turn blue, signalling that they can move back to the start position and begin the next trial. If neither criteria were met, they heard a loud beep and the start position turned red, signalling that they must return to the start position and repeat the trial until they satisfy both criteria. This feedback ensured participants did the task as well as they could, but these trials were not used in EEG analyses. One block of familiarization trials consisted of reaching to 12 target locations, which are the same targets they will experience in both the rotation and mirror perturbation types. Participants completed two familiarization trial blocks (24 trials total).

*Aligned trials*. Aligned trials followed the same procedure as familiarization, except participants advanced to subsequent trials, whether or not they satisfied the movement speed and endpoint position hold criteria. The start circle still turned blue or red to remind them of the speed and hold criteria. Each block included 12 target locations, with four blocks total (48 trials), providing baseline data for the perturbed trials.

*Random rotation trials*. Before each set of perturbation trials, we randomly rotated cursor feedback (±15°, ±25°, ±35°). Each rotation magnitude and direction appeared once in a shuffled manner before repeating, thereby making these rotations appear equally often but in an unpredictable manner. Each block included the corresponding six targets that will be encountered in the upcoming perturbation trials. Participants completed eight blocks (48 trials). These trials served to wash out prior learning before participants encountered the second perturbation, and as a control condition, since participants could not learn such unpredictable rotations but still experienced errors.

*Perturbed trials*. At the beginning of these trials, we informed participants that cursor feedback would be altered, and they should compensate for this. We perturbed visual feedback of the cursor in two ways. In fixed rotation trials, the cursor was rotated 30° counterclockwise relative to the hand position, and participants had to move 30° in the opposite direction to compensate. Although we intended to counterbalance the rotation direction to match movement paths between perturbation types (clockwise and counterclockwise, see Methods – Stimuli), a coding error restricted the perturbation in the counterclockwise direction. However, our prior work shows rotation direction does not affect adaptation (Gastrock et al., 2024), and we confirmed that despite this error, we do not find behavioral differences and our reported EEG effects remain robust (see R notebook, Gastrock et al., 2025). Thus, we included all rotation trials for data analysis. In mirror reversed trials, cursor feedback was flipped across either the vertical or horizontal midline axis, requiring movements to the opposite side of the mirror axis. Participants trained on one perturbation type before the other. Each block included six targets, and participants completed 15 blocks per perturbation type (90 trials) to ensure learning saturation.

### Behavioral data analysis

While not the focus of the current study, we investigated participants’ behavioral performance to confirm that we can replicate findings from our previous study that compared rotation adaptation with mirror reversal learning (Gastrock et al., 2024), and to inform trial categorization into the different sub-conditions for our EEG analyses (see EEG acquisition and preprocessing). As such, we do not perform an exhaustive set of statistical analyses for all behavioral results we present, and instead show figures containing the mean and 95% confidence intervals of each dependent variable we consider. We do, however, confirm that participants learned to compensate for the fixed rotation and mirror reversal perturbations using Bayesian t-tests that compare the amount of compensation across initial and later blocks of training. For these tests, we report Bayesian statistics to show evidence in support of the alternative hypothesis over the null hypothesis (i.e., BF10 values). We also report more detailed results from both frequentist and Bayesian tests in our accompanying R notebook (Gastrock et al., 2025). All behavioral data preprocessing and analyses were conducted in R version 4.2.2 (R Core Team, 2022).

### Amount of compensation

We quantified reach performance using the angular reach deviation of the hand, which is the angular difference between a straight line connecting the start position to the target and a straight line connecting the start position to the endpoint of the reach. Thus, the angular deviations for aligned reaches are expected to be close to zero. For both random rotation and perturbed trials, we first corrected for individual baseline biases by calculating the average angular deviation for each target within each participant during aligned trials, and subtracting this from angular deviations during random rotation or perturbed trials. Given that the fixed rotation perturbation was 30°, full compensation for this perturbation type corresponded to an angular reach deviation of 30°. Similarly, full compensation for the random rotation trials depended on the rotation magnitude and direction in each trial (±15°, ±25°, ±35°).

For the mirror perturbation however, compensation depended on how far the target was from the mirror axis. Thus, full compensation for the mirror perturbation could either be 15°, 30°, or 45°. Since full compensation differed across the perturbation types, we made compensation measures comparable by converting the angular reach deviation measures during aligned, random rotation, and perturbed trials into percentages of full compensation.

### Movement analyses

We also show how other measures related to the reaching movement, including reaction time, movement time, and path length, replicate findings from our previous study (Gastrock et al., 2024).

*Reaction time*. We defined reaction time (RT) as the time elapsed between the go signal onset and when the hand-cursor has moved 0.5 cm away from the center of the start position.

*Movement time*. We defined movement time (MT) as the interval between the first sample in which the hand cursor was more than 0.5 cm from the center of the start position and the first sample after the hand-cursor exceeded the distance to the center of the target position.

*Path length*. We defined path length (PL) as the total distance travelled between movement onset and offset (i.e., MT start and end), calculated from the x-y coordinates of the reach trajectory. The shortest path to the target is a straight line that starts from 0.5 cm away from the center start position, and ends at the target distance (shortest PL = 4.08 cm).

### EEG acquisition and preprocessing

EEG activity was recorded continuously at a sampling rate of 512 Hz, using a 64-channel ActiveTwo system (BioSemi), referenced to the CMS/DRL electrodes located between the Pz and POz electrodes, on the left and right side of the cap midline respectively. Electrodes were mounted on an elastic cap and placed according to the extended 10-20 EEG system. Each participant’s head circumference determined the cap size used, which consequently determined the electrode locations. The electrode offsets, which are voltage differences between the CMS and each active electrode, were maintained at ±25 µV.

### EEG preprocessing

EEG data were preprocessed and analyzed using MNE-Python version 1.2.1 (Gramfort et al., 2013). Continuous signals for each participant were re-referenced to the left and right mastoids, bandpass filtered between 0.1 and 200 Hz, with a notch filter at 60 Hz and its harmonics to remove line noise. We then segmented the signals into epochs from -2 to +2 seconds time-locked to the go signal onset and the feedback onset. For artifact removal, we applied a high pass filter at 1 Hz and subjected the data to an extended infomax Independent Component Analysis (ICA). We classified the different components using ICLabel (Pion-Tonachini et al., 2019), where components identified as muscle, eye, or channel noise with ≥ 80% probability were removed. We then applied the ICA results to the filtered data (i.e., subtracted identified artifacts from signal), before downsampling to 200 Hz.

As we were interested in EEG activity for time periods before and after the reaching movement, we time-locked to the go signal onset to investigate movement preparation, and time-locked to the feedback onset to investigate post-movement error processing. For each time-locked event, we wanted to test how EEG activity changed as one learned to compensate for the different perturbation types. That is, we expected larger errors to occur during the early stages of training, with errors becoming smaller as one progressed to the later stages of training. However, we did not observe such a pattern across the behavioral data of participants, as there was large variability in participant performance across trials especially for the mirror perturbation (Fig. 2A). We instead present two splits of the EEG data, one based on training phase (early vs. late), and one based on error size (large vs. small).

**Figure 2.**
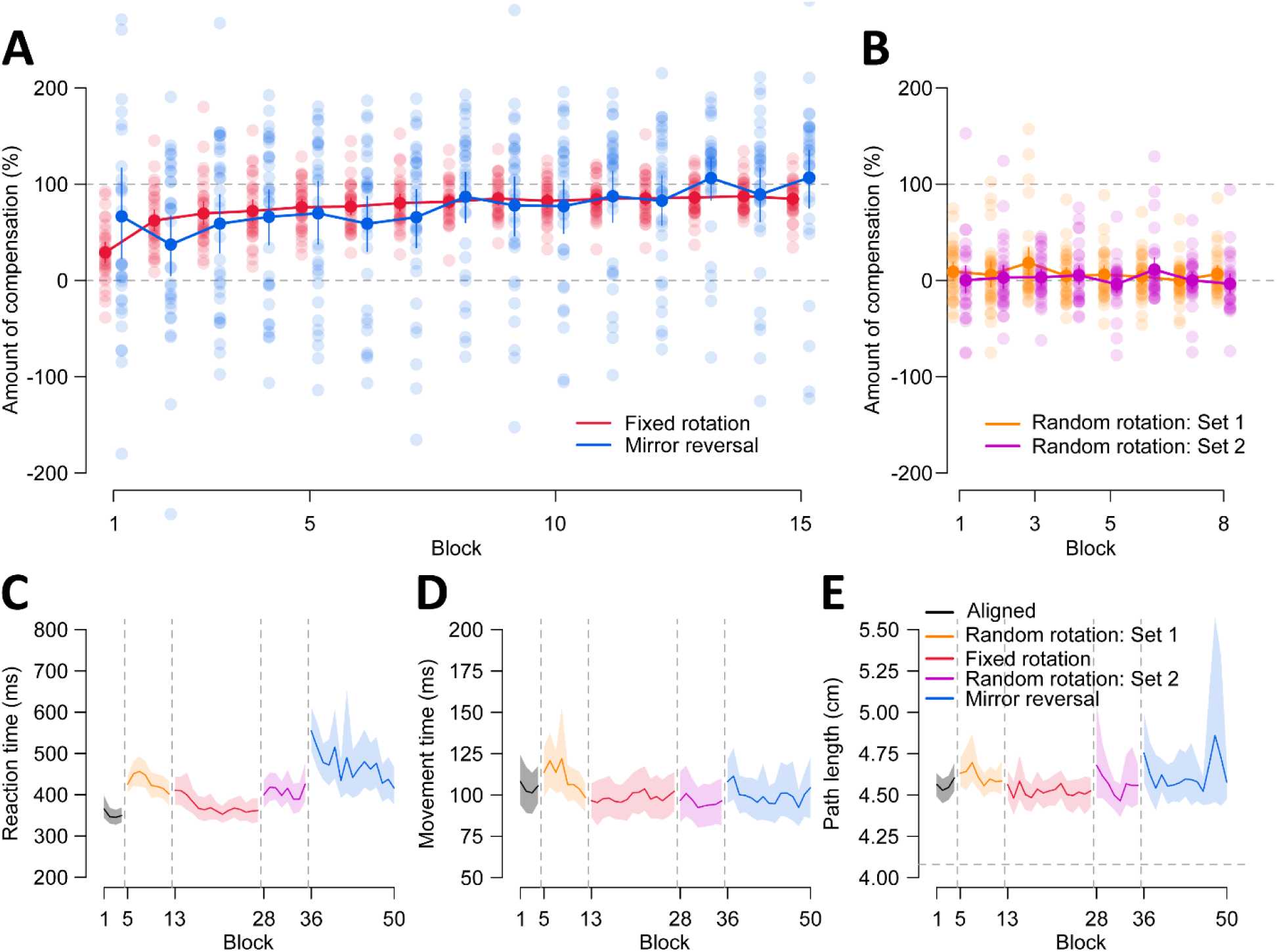
Reach performance and movement measures across trial types. To compare across trial types, angular reach deviations were converted to percentage of compensation. The grey dashed line at 100% indicates full compensation and 0% indicates no compensation (i.e., reaching directly to the target). Individual data are shown as faint dots, while means and 95% confidence intervals are shown with solid dots and error bars. **(A)** Participants learned to compensate for both perturbations, with greater variability for the mirror reversal. (**B**) "Set 1" and "Set 2" refer to the first and second sets of random rotation trials preceding each perturbation training respectively. As expected, no learning occurred for random rotations. (**C**) Reaction time (RT) is the time elapsed between the go signal onset and when the hand-cursor has moved 0.5 cm away from the center of the start position. RTs were slower for all trial types compared to the aligned trials. (**D**) Movement time (MT) is the time elapsed between the first sample when the hand-cursor is >0.5 cm away from the center of the start position and the first sample when the hand-cursor is greater than the distance to the center of the target. Overall, there are no substantial differences in MTs across trial types. (**E**) Path length (PL) refers to the total distance travelled between movement onset and offset (i.e., MT start and end), calculated from the x-y coordinates of the reach trajectory. The shortest path to the target is a straight line that starts from 0.5 cm away from the center start position, and ends at the target distance (4.08 cm). Overall, there are no differences in path length across trial types. For (**C-E**), each block is an average across six trials (12 trials for aligned). Solid lines and shaded regions are means and 95% confidence intervals across participants.

*Training phase: Early vs. Late*. Given the quick progression of learning that we observed for both the fixed rotation and mirror perturbations (Fig. 2A), we used the first two blocks of trials (first 12 training trials) from each participant to define early training in both perturbations. The amount of compensation plateaus for the rest of training, so we used the last six blocks of trials (last 36 training trials) for each participant to define late training in both perturbations. The random rotation trial type is shorter and occur before each perturbation training. For these, we used the first two blocks of trials (Random rotation: Set 1; Fig. 1B & 2B) to define early training and the last four blocks of trials (Random rotation: Set 2; Fig. 1B & 2B) to define late training.

*Error size: Large vs. Small*. The error magnitude experienced within a given trial is dependent on the perturbation type experienced. For the fixed rotation, we sorted all errors for each participant. We defined small errors as the 36 lowest values and large errors as the 18 highest. This mirrors the data quantity used in the early (12 trials)/late (36 trials) split, but we included 18 rather than 12 of the largest errors, as they remain distinct from small errors while allowing a modest increase in power. For the mirror perturbation, we first identified errors relative to each target location, as each target produced a different error magnitude. We then sorted these relative error sizes for each participant, before defining small (36 trials) and large (18 trials) errors similarly to the fixed rotation. For the random rotation trials, we combined all random rotation trials that occurred prior to each perturbation type. We then identified errors relative to each target location, before sorting these relative errors and defining which trials belonged with the small (36 trials) and large (18 trials) error conditions.

For both training phase and error size conditions, we used all 48 aligned trials as baseline data to compare with the perturbed trials.

### EEG time domain analysis

We compare event related potentials (ERPs) generated from the training phase and error size conditions both prior to and following the reaching movement.

### EEG data comparison steps

The comparisons across the different training phase and error size conditions are implemented in three steps. First, we compare whether signals for each of the four conditions (early, late, small, or large) in the random rotation, fixed rotation, and mirror reversal perturbations are different from signals in the baseline aligned trials. Second, we calculate difference waves between each condition and the aligned condition, such that we can compare training phase and error size conditions directly (early vs. late, large vs. small). Third, we calculate difference waves between training phase (late minus early) and error size (small minus large) conditions in each perturbation type, such that we can compare across perturbation types.

### Movement preparation

Prior to the reaching movement in each trial, participants were cued of the upcoming target location for one second, before receiving the go signal to move towards the target. Therefore, we time-locked to the go signal onset, used 1300 - 1000 ms prior the go signal onset as baseline, and implemented our outlined comparison steps during the one second preceding the go signal. For these comparisons, we included six electrodes located around the scalp midline and motor areas to investigate the readiness potential (RP; FCz, Fz, C3, Cz, C4, Pz; as in Reznik et al., 2018; Wen et al., 2018). While the RP has been shown to reflect effector-specific preparatory activity, called the Lateralized Readiness Potential (LRP; Smid et al., 1987; de Jong et al., 1988; Gratton et al., 1988), we observed that our measured RPs seem to take into account the planned movement direction on the workspace. That is, C3 had a more negative RP than C4 when the upcoming movement was to the right side of the workspace, and vice-versa for leftward movements, despite all reaches being performed with the right hand. Notably, this effect is observed only for the targets along the horizontal axis of the workspace. As such, we calculated LRPs for epochs with these target locations using the equation (de Jong et al., 1988; Luck & Kappenman, 2013): LRP = (right movement C3 – right movement C4) – (left movement C3 – left movement C4) We then used these calculated LRPs to compare across conditions.

### Post-movement error processing

After participants completed the reaching movement within a trial, they received visual feedback of where the cursor ended relative to the target for one second. Thus, we time-locked EEG activity to the feedback onset, used 100 ms prior to the feedback onset as baseline, and implemented our outlined comparison steps on the signals during the one second following feedback onset. For these comparisons, we included 10 fronto-central-parietal electrodes (F3, Fz, F4, FCz, C3, Cz, C4, P3, Pz, P4; as in (Maurer et al., 2019), to investigate the Feedback Related Negativity (FRN) and subsequent P3 components post-feedback.

### EEG time-frequency analysis

We transformed the preprocessed epochs in the time-frequency domain using Morlet wavelet convolution (6 cycles, with log-spaced frequencies ranging from 6 to 35 Hz) for the full epoch for each trial (2 seconds before go signal to 2 seconds after feedback onset) but only analyzed power for the one second preceding the go signal onset and following the feedback onset. Explicit and implicit learning are expected to contribute differently to fixed rotation and mirror reversal perturbations, given that the mirror perturbation is more cognitively demanding while compensation for the relatively small rotation is mostly unconcscious (Gastrock et al., 2020). Thus, we analyzed pre-specified regions of interest (ROIs; as in (Jahani et al., 2020) sensitive to these processes. The medial frontal electrodes (F1, Fz, F2, FC1, FCz, FC2, C1, Cz, C2) were chosen for their link to explicit adaptation, and lateral central electrodes contralateral to the right hand (C5, C3, CP5, CP3, CP1, P5, P3, P1) for their link to implicit adaptation.

For each ROI, we computed mean power across three frequency bands: theta (6–8 Hz), alpha (9–13 Hz), and beta (13–25 Hz), each previously shown to change during adaptation tasks (e.g., Gentili et al., 2011; Tan et al., 2014). We then compared average power across conditions using the same procedures described above.

### EEG statistical analysis

For each of the comparison steps we implemented, we compare whether two signals are statistically different from each other using cluster-based permutation t-tests (Maris & Oostenveld, 2007; Sassenhagen & Draschkow, 2019). This method allows signal comparisons across multiple timepoints, while inherently controlling for multiplicity. Instead of testing each timepoint independently, the method first identifies clusters of consecutive timepoints in which t-values exceed a predefined threshold (p = 0.05). These timepoints are grouped into clusters based on temporal adjacency. To determine whether a cluster is statistically significant, we then compare its cluster-level statistic (the sum of t-values within the cluster) against a distribution of cluster statistics generated with 1,000 permutations (with replacement) of the data. A cluster is considered significant (p < 0.05) only if its statistic falls within the extreme tail of this permutation distribution, indicating that such a cluster is unlikely to arise by chance. Both identified and significant clusters are displayed at the bottom of the EEG figures and their corresponding p-values are reported in the Results section.

### Code accessibility

Behavioral and EEG data, along with analyses scripts are available on Open Science Framework (Gastrock et al., 2025: <u>10.17605/OSF.IO/Q5N6J</u>).

## Results

We first confirm that participants successfully learn to compensate for both the fixed rotation and mirror reversal perturbations, consistent with findings from our previous study (Gastrock et al., 2024), and that no learning progresses during random rotation trials. As expected, participants learn both fixed rotation and mirror reversal perturbation types quickly, with learning in the mirror perturbation showing much larger inter-participant variability (Fig. 2A). Using a paired Bayesian t-test, we find evidence that the first block of fixed rotation trials differs from the last block of fixed rotation trials (BF > 1·10^8^), suggesting that participants compensated for the perturbation. For the mirror reversal perturbation, we find no evidence of a difference between the first and last blocks of training (BF = 0.447). However, this is likely due to the large inter-participant variability affecting the mean, as we see more learning progression occurring from the second block to the last block of training (Fig. 2A; BF = 13.120). We also observe that when encountering different error magnitudes and directions per trial in the random rotation blocks, participants do not learn to compensate (Fig. 2B; Random – Set 1, block 1 vs. last block: BF = 0.209; Random – Set 2, block 1 vs. last block: BF = 0.207). These findings confirm that participants are able to compensate for the rotation and mirror reversal perturbations, but not for the unpredictable random rotations.

We also investigate patterns of different movement measures across trial types. For reaction time (RT), we find that movement initiation slows down across all trial types compared to aligned baseline reaches, with RTs in the mirror perturbation being much slower than other trial types (Fig. 2C). For both movement time and path length measures, we generally do not observe differences across trial types (Fig. 2D-2E). These patterns of results are expected and similar to our previous findings (Gastrock et al., 2024). Thus, we use these behavioral results to inform how we categorize and investigate the corresponding EEG data.

### Extent of training and error magnitude modulate EEG activity across different perturbation types during pre- and post-movement

We first investigate EEG dynamics during perturbation training in the temporal domain. For movement preparation, we used permutation-based t-tests to compare the Readiness Potentials (RPs) between training phases and error sizes within each perturbation type, and across the different perturbation types, during the preparatory period prior to the go signal onset. For training phase comparisons, we find that late training in the fixed rotation exhibits a more positive-going RP just prior to the go signal onset, whereas there seems to not be any RP in early training (Fig. 3A, -0.15 – 0.00 s, p = 0.036). However, we do not observe such differences for the random rotation and mirror perturbations (i.e., no significant clusters; Fig. 3B-3C). The RPs in the fixed rotation hovers near zero early in training before becoming more positive later in training. While not statistically significant, the random rotation in contrast shows a positive deflection throughout, with early training being more positive than late. This flip in the relationship is evident with the difference we find when comparing fixed and random rotation perturbations (Fig. 3D, -0.18 – 0.00 s, p = 0.031), suggesting that preparatory activity is modulated by training phase differently for fixed and random rotations. For error size comparisons, we do not find any differences between small and large errors within and across the different perturbation types (Fig. 3E-3H). Taken together, these results suggest that the phase of training modulates preparatory EEG activity specifically for the fixed rotation perturbation, whereas the magnitude of the upcoming movement error is not influenced by movement preparation.

**Figure 3.**
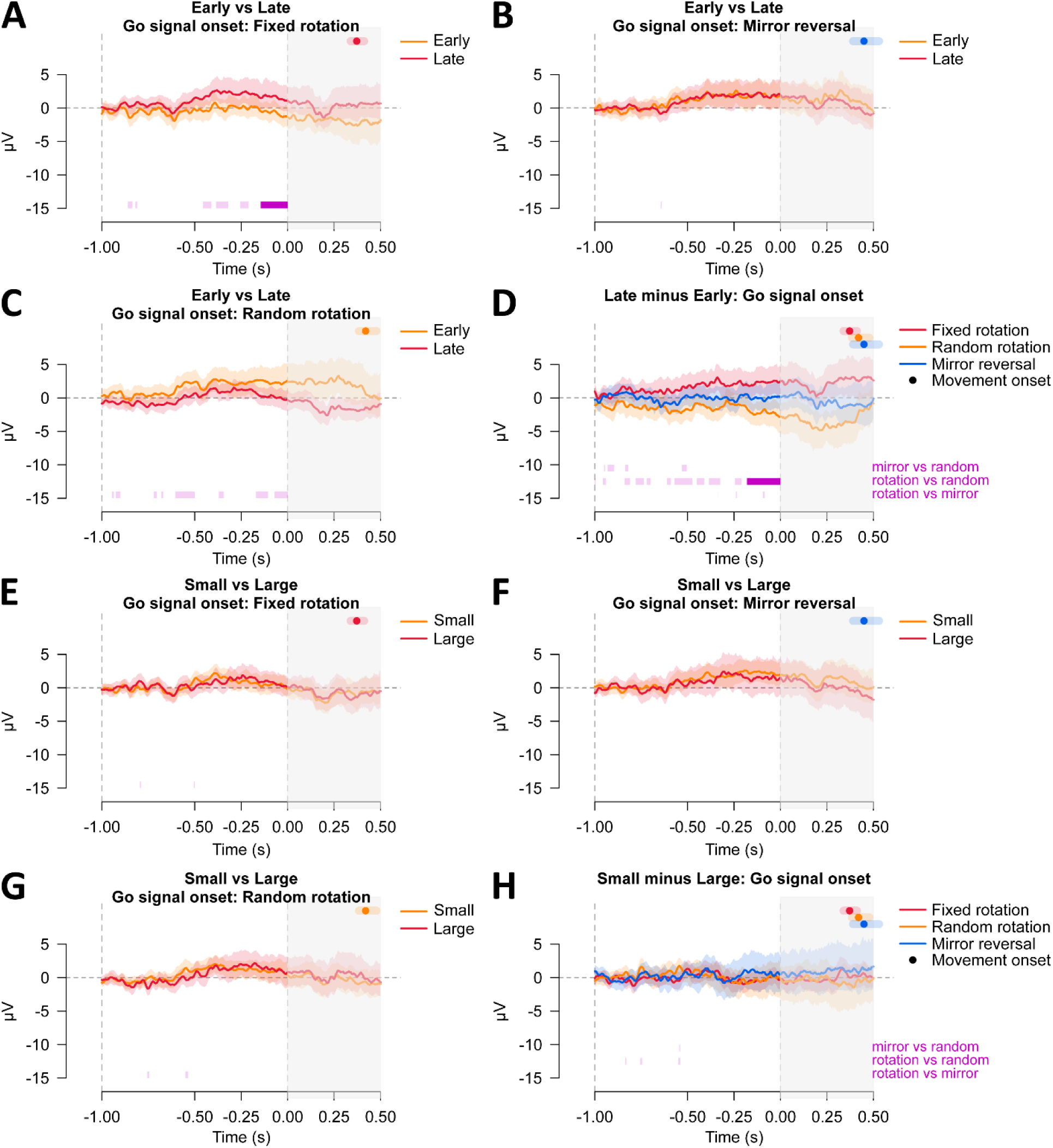
Event Related Potentials across perturbation types prior to the go signal onset. Within each perturbation type (fixed rotation, mirror reversal, random rotation), we compare early versus late training epochs (**A- C**) and epochs with small versus large errors (**E-G**), before calculating difference waves to compare across perturbation types for training phase (**D**) and error size (**H**) respectively. EEG activity is time-locked to the go signal onset. Solid lines and shaded regions show the signal means and 95% confidence intervals (CIs) across participants for the second preceding the go signal onset. Solid dots and error bars after the go signal onset show movement onset means and 95% CIs for each specific perturbation type. Light-shaded purple bars at the bottom of each panel show identified clusters from the permutation-based t-tests to compare the two signals, and solid purple bars show statistically significant clusters.

Although participants consistently used their right hand to perform the reaching movements, we still observed Lateralized Readiness Potentials (i.e., C3 electrode has a more negative RP than the C4 electrode when the upcoming movement was to the right side of the workspace, and vice-versa for leftward movements). Importantly, we find LRPs across all trial types (Fig. 4). However, these LRPs are neither modulated by training phase nor error magnitude (R notebook, Gastrock et al., 2025). Taken together, these results suggest that LRPs not only process effector-specific preparatory activity, but also accounts for the planned movement direction.

**Figure 4.**
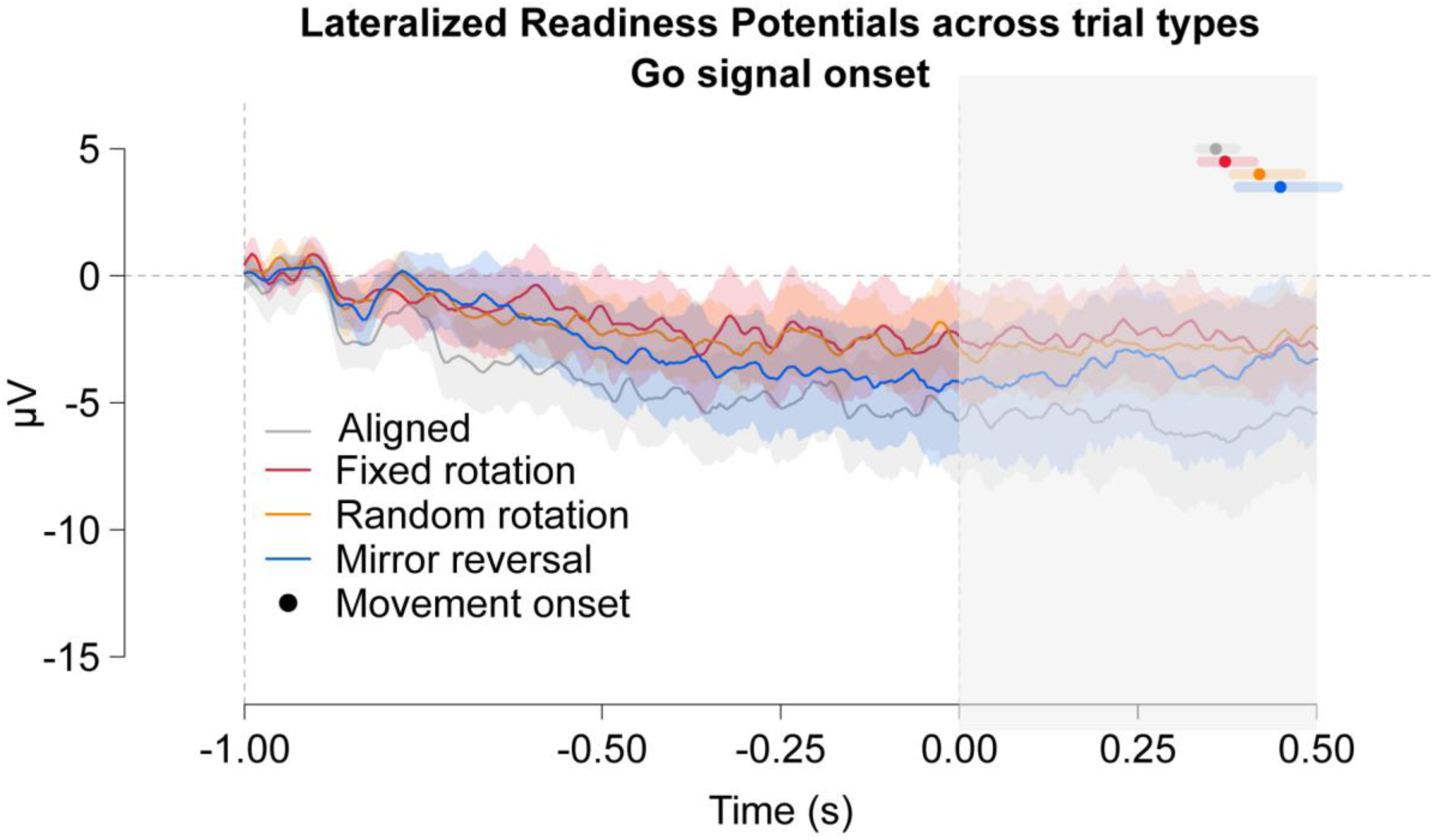
Lateralized Readiness Potentials across trial types during movement preparation. For movements to targets on the left or right side of the workspace horizontal axis, we calculated Lateralized Readiness Potentials (LRPs) using electrodes C3 and C4. We show these LRPs across the different trial types: aligned, fixed rotation, random rotation, and mirror reversal. Solid lines and shaded regions indicate LRP signal means and 95% confidence intervals (CIs) across participants for the second preceding the go signal onset. Solid dots and shaded regions after the go signal onset show means and 95% confidence intervals of the movement onset for each trial type.

We then investigated error processing following a reaching movement, to assess the Feedback Related Negativity (FRN) and subsequent P3 components. As such, we used permutation-based t-tests to compare between training phases within each perturbation type, and across the different perturbation types, during the second that follows the feedback onset (Fig. 5A-5C). We observe a positive-going deflection, likely reflecting the P3 component given the timepoint at which the component’s peak occurs. For training phase, we find that early training exhibits more positive-going signal post-feedback compared to late training in both the fixed rotation (Fig. 5A, 0.05 – 0.34 s, p = 0.010) and random rotation (Fig. 5C, 0.17 – 0.54 s, p = 0.006), but not the mirror perturbation (i.e., no identified clusters; Fig. 5B).

**Figure 5.**
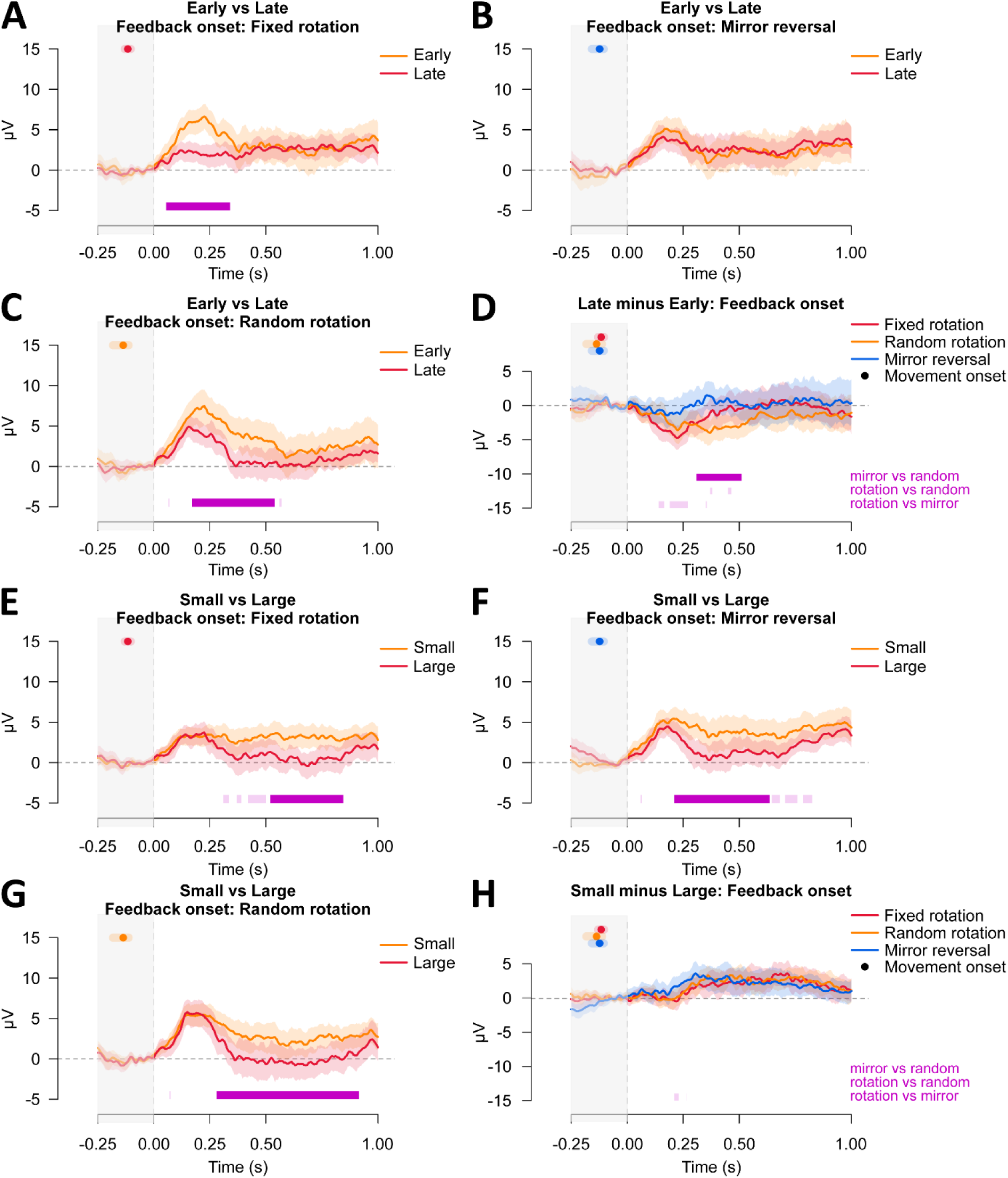
Event Related Potentials across perturbation types following post-feedback onset. Within each perturbation type (fixed rotation, mirror reversal, random rotation), we compare early versus late training epochs (**A-C**) and epochs with small versus large errors (**E-G**), before calculating difference waves to compare across perturbation types for training phase (**D**) and error size (**H**) respectively. EEG activity is time-locked to the feedback onset. Solid lines and shaded regions show the signal means and 95% confidence intervals (CIs) across participants for the second following the feedback onset. Solid dots and error bars prior to feedback onset show movement onset means and 95% CIs for each specific perturbation type. Light-shaded purple bars at the bottom of each panel show identified clusters from the permutation-based t-tests to compare the two signals, and solid purple bars show statistically significant clusters.

We then calculate difference waves between early and late training, to directly compare across perturbations, and find that the random rotation exhibits a more negative-going signal compared to the mirror perturbation (Fig. 5D, 0.31 – 0.51 s, p = 0.031). These results suggest a shift in error processing as learning progresses for rotation-type perturbations, which does not occur for the mirror perturbation.

We also compared small and large errors within and across perturbation types post-feedback onset (Fig. 5E-5G). We find that small errors consistently exhibit a more positive-going signal than large errors post-feedback (fixed rotation: 0.52 – 0.85 s, p = 0.007; mirror reversal: 0.21 – 0.64, p = 0.007; random rotation: 0.28 – 0.92s, p = 0.002). However, this small-large difference is about the same magnitude when comparing across perturbation types (Fig. 5H), as we find no significant clusters for these comparisons.

Thus, while error magnitude modulates EEG activity post-feedback, the processing of error magnitude does not differ across perturbation types.

### Extent of training and error magnitude modulate beta and alpha band power across different perturbation types during pre- and post-movement

In addition to our investigation of EEG activity along the temporal domain, we also explore how EEG dynamics in the frequency domain may change during learning. For these analyses, we focus on two specific regions of interest (ROIs; as in Jahani et al., 2020): the medial frontal area which has been shown to process more cognitive aspects such as explicit contributions to learning, and the lateral central area contralateral to the moving right hand which has been shown to process more unconscious or implicit contributions to learning. We then calculate the mean power within each participant for three different frequency bands: theta (6–8 Hz), alpha (9–13 Hz), and beta (13–25 Hz). We selected these frequency bands because previous studies have shown they synchronize and/or desynchronize during pre- and post-movement in adaptation tasks (Gentili et al., 2011; Tan et al., 2014). Here, we examine how these frequency changes may differ across perturbation types. While detailed comparisons are found in our R notebook (Gastrock et al., 2025), we only present comparisons across perturbation types here, to keep the discussion concise.

We first investigate whether training phase and error size differences across perturbation types, modulate the beta frequency band differently across the two ROIs during movement preparation and feedback processing. We know from previous research that pre-movement beta power in a subsequent trial decreases after encountering movement errors, with stronger attenuation for large errors early in learning than for smaller errors later in adaptation (Reuter et al., 2022). Similarly, post-movement beta is also more strongly attenuated following larger compared to smaller errors. Here, we examine how beta power during movement preparation relates to the upcoming movement error within that trial, as well as how post-movement beta is affected by such experienced error. Furthermore, we expect that reduced beta power in the lateral central area to represent sensorimotor adaptive (implicit) mechanisms, while reduced beta power in the medial frontal area to represent more deliberate (explicit) and strategic behavioral adjustments (Jahani et al., 2020). When comparing late minus early training differences across perturbations (Fig. 6A-6D), we only find differences in the medial frontal area post-feedback onset. In figure 6B, the random rotation perturbation is in the positive direction compared to the other two perturbation types, and differs significantly from the fixed rotation (0.20 – 0.47 s, p = 0.016). Indeed, if we compare the early and late signals for the random rotation (R notebook, Gastrock et al., 2025), we find that beta power is attenuated (more negative) during early training and increases back to baseline levels (zero) during late training, suggesting a change in error-processing that is specific to the random rotation perturbation.

**Figure 6.**
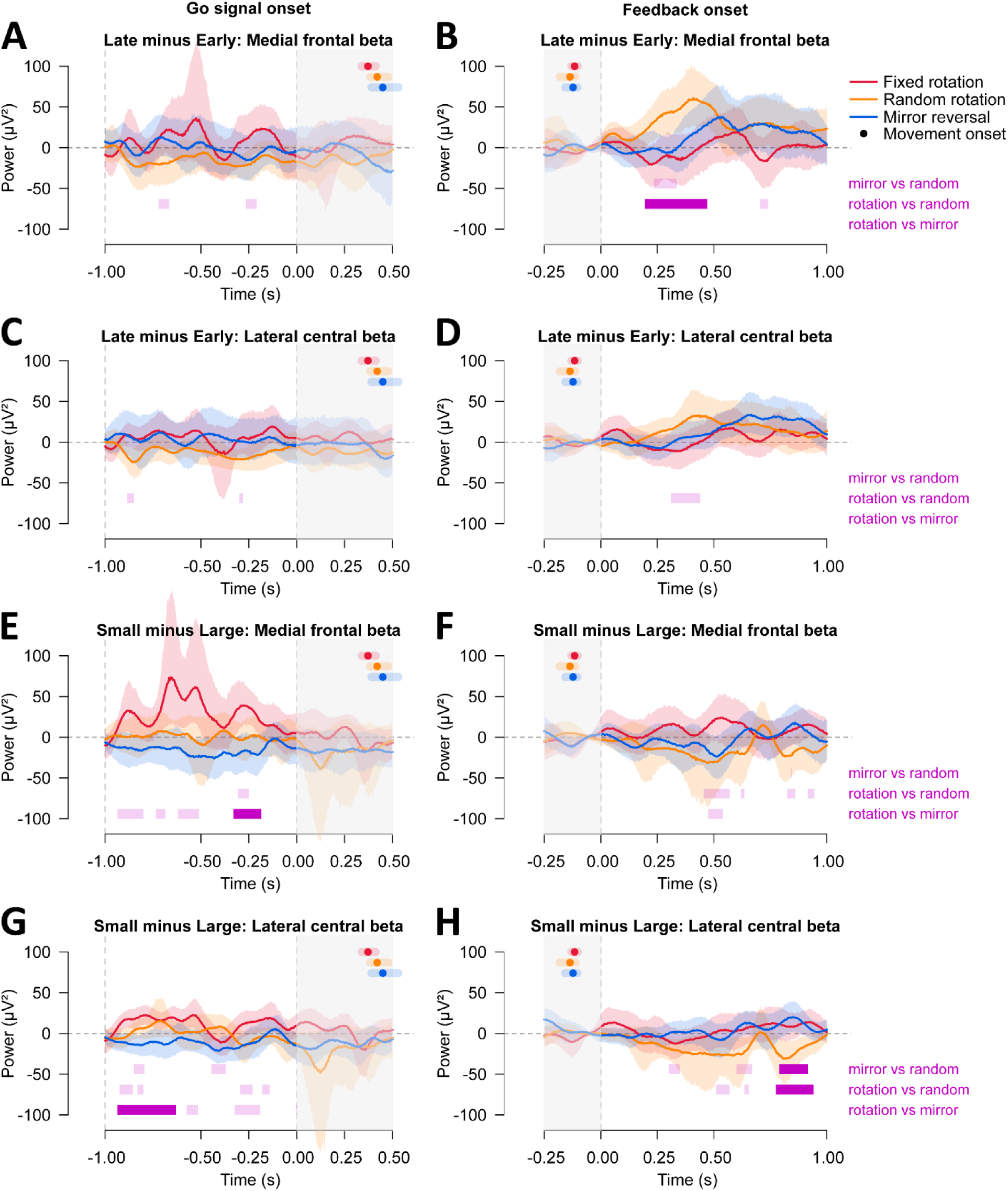
Changes in EEG power in the beta band during movement preparation and feedback processing. We show EEG power in medial frontal and lateral central areas that correspond to late minus early training differences (**A-D**) and small minus large error differences (**E-H**) across the different perturbation types. The left column contains EEG activity time-locked to the go signal onset, while the right column shows EEG activity time-locked to the feedback onset, to investigate movement preparation and feedback processing, respectively. Solid lines and shaded regions show the signal means and 95% confidence intervals (CIs) across participants, color-coded according to each perturbation type. Light-shaded purple bars at the bottom of each panel show identified clusters from the permutation-based t-tests to compare the two signals, and solid purple bars show statistically significant clusters. In all panels, Solid dots and shaded regions after the go signal onset or before the feedback onset show means and 95% confidence intervals of the movement onset for each perturbation type.

When comparing small minus large error differences across perturbations (Fig. 6E-6H), we find differences in the medial frontal area between the fixed rotation and mirror perturbations during movement preparation (Fig. 6E, -0.33 - -0.19 s, p = 0.038), as well as in the lateral central area (Fig. 6G, - 0.94 - -0.63 s, p = 0.007). In both comparisons, the fixed rotation signal is in the positive direction, where we observe that beta power is attenuated (more negative) for larger errors, and increases back to baseline levels for small errors. Thus, attenuated beta power during movement preparation is associated with larger errors in the upcoming movement. Finally, in the lateral central area post-feedback onset (Fig. 6H), we find that the random rotation differs from both the fixed rotation (0.78 – 0.94 s, p = 0.030) and mirror perturbations (0.79 – 0.92 s, p = 0.036). In this case, the random rotation is in the negative direction, where beta power is increased (more positive) following large errors but decreases to baseline levels following small errors. Taken together, it seems that beta power is modulated differently across the medial frontal and lateral central areas following feedback in the random rotation perturbation, while beta power during movement preparation influences error magnitude differently for the fixed rotation and mirror perturbations.

We then investigate how the different perturbation types affect alpha frequency band activity across different parts of the task. Frontal alpha frequency band power during movement preparation is expected to be positively associated with motor performance (Gentili et al., 2011). Specifically, alpha power desynchronizes during early stages of training but resynchronizes as learning progresses (i.e., progressive idling). Interestingly, alpha synchronization is also observed in temporal and parietal areas as adaptation progresses (Gentili et al., 2011). Thus, alpha synchronization seems to be related to updating internal models that allows one to adapt to a given perturbation. In the current study, we do not observe any alpha power modulations that differed across perturbation types when comparing early versus late training (Fig. 7A–7D), or between small and large error magnitudes (Fig. 7E, 7F, 7H), during either movement preparation or feedback processing. However, one exception we find is that there is a relationship between preparatory alpha power in the lateral central area and the size of the subsequent error, that is different for fixed rotation and mirror perturbations (Fig. 7G, -0.97 - -0.68 s, p = 0.023).

**Figure 7.**
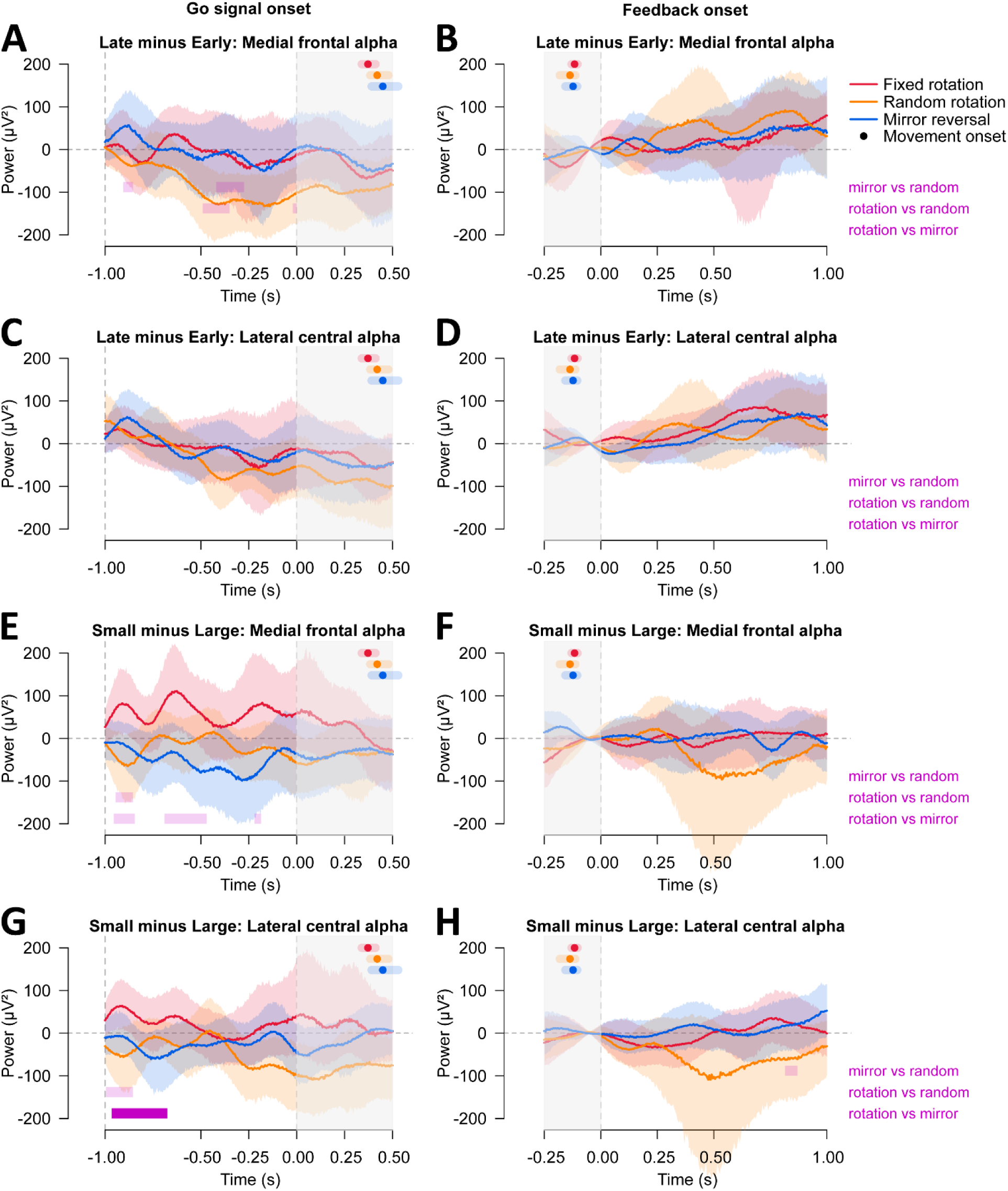
Changes in EEG power in the alpha band during movement preparation and feedback processing. We show EEG power in medial frontal and lateral central areas that correspond to late minus early training differences (**A-D**) and small minus large error differences (**E-H**) across the different perturbation types. The left column contains EEG activity time-locked to the go signal onset, while the right column shows EEG activity time-locked to the feedback onset, to investigate movement preparation and feedback processing, respectively. Solid lines and shaded regions show the signal means and 95% confidence intervals (CIs) across participants, color-coded according to each perturbation type. Light-shaded purple bars at the bottom of each panel show identified clusters from the permutation-based t-tests to compare the two signals, and solid purple bars show statistically significant clusters. In all panels, Solid dots and shaded regions after the go signal onset or before the feedback onset show means and 95% confidence intervals of the movement onset for each perturbation type.

Given that the difference between small and large errors for the fixed rotation is in the positive direction (Fig. 7G), this suggests that preparatory alpha synchronizes for movements that produce smaller compared to larger error magnitudes. Although we observe a cluster of timepoints that show this relationship, we note that these error size comparisons in alpha power for the fixed rotation (large vs. small) are not statistically significant (see R notebook, Gastrock et al., 2025). In contrast, we do not find the same synchronization patterns for the mirror reversal or random rotation perturbations. Thus, we only find evidence for lateral central alpha synchronization during movement preparation in the fixed rotation perturbation.

Finally, we investigate how the different perturbation types affect theta frequency band activity across different parts of the task. We expect motor performance to be positively associated with frontal theta power during movement preparation (Gentili et al., 2011). That is, theta power in the frontal electrodes during movement preparation should increase as one learns to compensate for the perturbation. However, we find that theta power is not modulated by either training phase or error magnitude during movement preparation in the medial frontal area. Importantly, this lack of a difference seems to be consistent across perturbation types, as we do not observe differences across the fixed rotation, mirror reversal, and random rotation perturbations (Fig. 8A, 8E). We also explore whether theta power is modulated post-feedback (Fig. 8 right column) and in the lateral central area (Fig. 8C-8D, 8G-8H), and find no differences across perturbation types as well. Thus, we find no overall theta power differences across perturbation types both during movement preparation and post-movement feedback processing.

**Figure 8.**
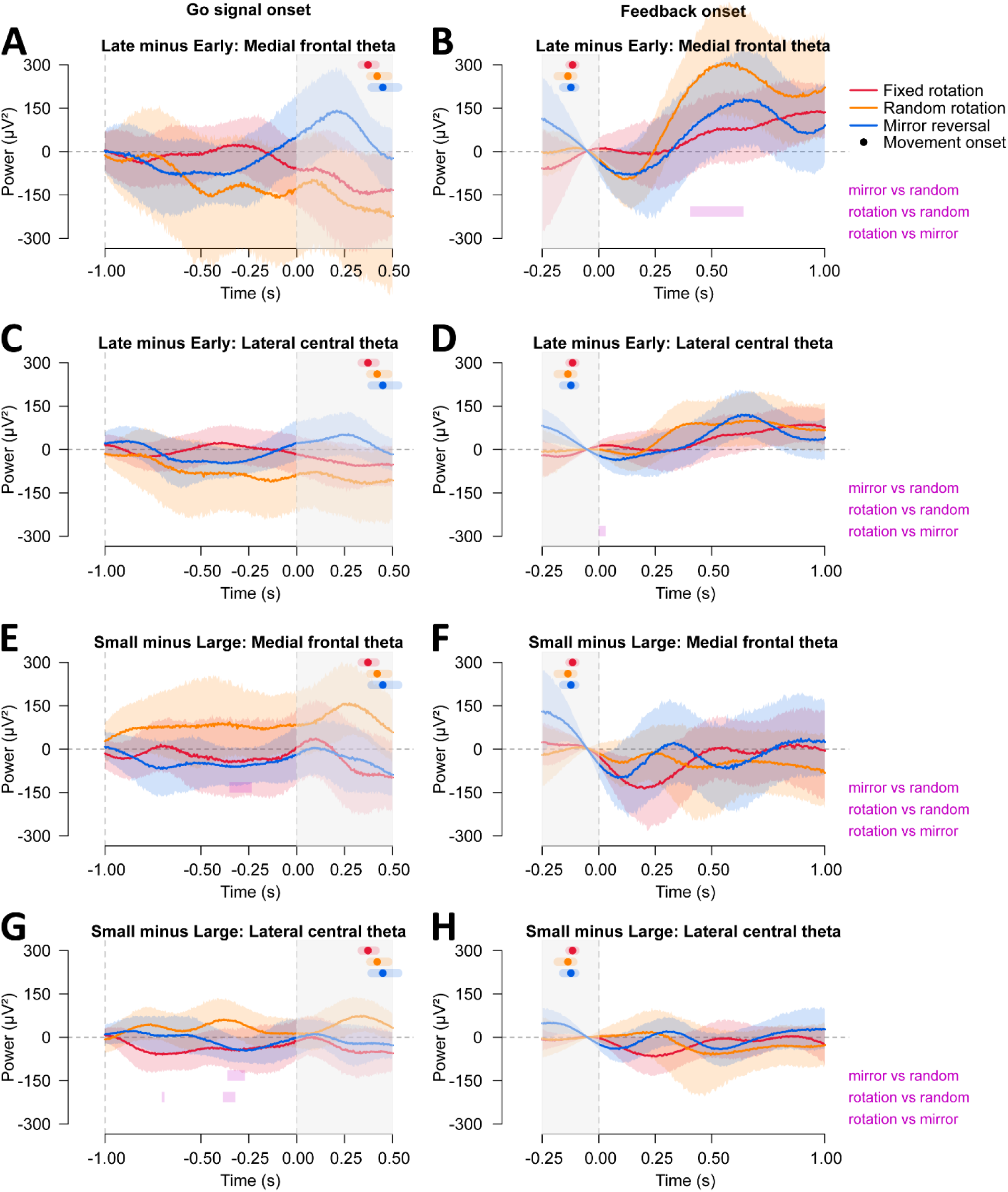
Changes in EEG power in the theta band during movement preparation and feedback processing. We show EEG power in medial frontal and lateral central areas that correspond to late minus early training differences (**A-D**) and small minus large error differences (**E-H**) across the different perturbation types. The left column contains EEG activity time-locked to the go signal onset, while the right column shows EEG activity time-locked to the feedback onset, to investigate movement preparation and feedback processing, respectively. Solid lines and shaded regions show the signal means and 95% confidence intervals (CIs) across participants, color-coded according to each perturbation type. Light-shaded purple bars at the bottom of each panel show identified clusters from the permutation-based t-tests to compare the two signals, and solid purple bars show statistically significant clusters. In all panels, Solid dots and shaded regions after the go signal onset or before the feedback onset show means and 95% confidence intervals of the movement onset for each perturbation type.

## Discussion

The current exploratory study investigates how the brain prepares movements and processes errors during both visuomotor rotation adaptation and mirror reversal de novo learning. We used EEG recordings during movement preparation and post-movement feedback to assess how perturbation type, training phase (early vs. late), and error size (large vs. small) shape neural dynamics. Behaviorally, participants successfully compensate for both the fixed rotation and mirror reversal but not the random rotation, with greater variability and slower reaction times during mirror reversal training. Aftereffects are only observed following fixed rotation training, consistent with our prior findings (Gastrock et al., 2024).

During movement preparation (see Table 1), we observe that 1) fixed and random rotations modulate the RP in opposite ways across training (the fixed rotation becomes more positive late in training, whereas the random rotation is more positive early in training), 2) LRPs reflect processing of the planned movement direction, and 3) error size differentially influences beta and alpha power across perturbations: beta attenuation for large-error movements in medial and lateral central areas, and alpha synchronization for small-error movements in the lateral central area, are both observed more clearly in the fixed rotation than in the mirror reversal. During feedback processing (Table 1), 1) training phase modulates the P3 for both fixed and random rotations, with a clearer early-versus-late difference in the fixed rotation, and 2) small errors elicit a sustained positivity across perturbations. In the frequency domain, 3) the random rotation shows attenuated medial frontal beta during early training, and increased lateral central beta power following large errors. Together, these results show that the brain engages different preparatory and error-processing mechanisms across different types of motor learning.

**Table 1.** Summary of EEG effects. We summarize the EEG components where we observed significant effects during both movement preparation and feedback processing. For each component, we describe its expected functional role based on prior research and report the effects observed in our study, noting the perturbation types in which they occurred.

| EEG component | Expected functionality | Perturbation type | Observed effect |
| --- | --- | --- | --- |
| **movement preparation** |  |  |  |
| RP | movement preparation; increased negativity with expected sensory consequences | fixed rotation | increased positivity from early to late training |
| LRP | effector-specific preparation | all | preparation for intended movement direction |
| Beta (lateral central) | resynchronization with improved motor performance; linked to implicit learning | fixed rotation | resynchronization with small errors |
| Beta (medial frontal) | resynchronization with improved motor performance; linked to explicit learning | fixed rotation | resynchronization with small errors |
| Alpha (lateral central) | resynchronization with improved motor performance; task engagement | fixed rotation | resynchronization with small errors |
| **feedback processing** |  |  |  |
| P3 | cognition and attention; scales with error size | fixed and random rotation | amplitude decreases from early to late training |
| SLP | negative association with explicit learning | all | sustained positivity with small errors |
| Beta (lateral central) | resynchronization with improved motor performance | random rotation | increased with large vs. small errors |
| Beta (medial frontal) | resynchronization with improved motor performance | random rotation | resynchronization from early to late learning |

Our findings show that movement preparation evolves differently across learning in the different perturbation types. In the temporal domain, all perturbations elicited the expected RP negativity (Kornhuber & Deecke, 1965; Reznik et al., 2018), confirming its role in preparatory activity. However, RP modulation across training differed depending on perturbation. In the fixed rotation, the RP became more positive from early to late learning, consistent with a shift toward using an updated internal model as sensory prediction errors decrease. This interpretation aligns with prior work linking the RP to expectations about sensory consequences of actions, even if these previous studies used only button presses for movements (Jo et al., 2014; Reznik et al., 2018; Vercillo et al., 2018; Wen et al., 2018; Pinheiro et al., 2020). In contrast, the mirror reversal showed no RP differences across training, consistent with its reliance on more explicit processes rather than implicit sensory prediction error–based learning, and potentially related to observed behavioral measures of slower reaction times and greater inter-participant variability (Gastrock et al., 2024). Although the random rotation did not exhibit a significant early-late RP difference, its pattern was opposite that of the fixed rotation, possibly reflecting initial attempts to adapt followed by disengagement once the perturbation was recognized as unpredictable. For the LRP, prior work identifies it as an effector-specific preparatory component (Smid et al., 1987; de Jong et al., 1988; Gratton et al., 1988; Reznik et al., 2018). Despite participants always using the right hand, we observed LRPs driven by movement direction (leftward vs. rightward reaches) rather than perturbation type, indicating that LRPs also reflect movement direction planning. Taken together, RP changes across training appear to reflect updates to expected sensory consequences during motor adaptation, whereas LRPs reflect the intended movement direction.

In the frequency domain, we find that both beta and alpha activity vary with the magnitude of the upcoming movement error. For beta activity, fixed rotation training shows distinct patterns from mirror reversal training, across both the medial frontal and lateral central regions. During fixed rotation training, beta power is attenuated before large error movements and returns toward baseline for small error movements, consistent with prior adaptation work showing beta desynchronization early in learning and resynchronization as errors decrease (Tan et al., 2014; Ozdenizci et al., 2017). Notably, beta modulation during fixed rotation training emerged in both regions of interest. Previous research has shown that lateral central beta attenuation is linked to implicit adaptation, while medial frontal beta attenuation has been associated with explicit re-aiming (Jahani et al., 2020). Although we did not measure aiming strategies directly, changes in neural dynamics in both regions during fixed rotation training are likely due to both implicit and explicit processes contributing to motor adaptation (Mazzoni & Krakauer, 2006; Taylor et al., 2010; Werner et al., 2015; Modchalingam et al., 2019; Gastrock et al., 2020). For alpha activity, fixed rotation training shows a different pattern from mirror reversal training in the lateral central region. We observe a small, non-significant cluster in the fixed rotation condition in which alpha synchronization increases for small relative to large errors, a pattern not present in mirror reversal training. Increased alpha synchronization with smaller errors is often interpreted as reduced task engagement or cortical idling, typically seen in frontal but also temporal and parietal regions (Gentili et al., 2011). Thus, we find a modest indication of cortical idling around central and parietal electrodes contralateral to the moving hand (lateral central area) during fixed rotation training, though this should be interpreted cautiously given the small effect and contrasting evidence that contralateral motor alpha often decreases as performance improves (Desrochers et al., 2020). In contrast, neither beta nor alpha activity during mirror reversal training differentiates small from large errors. This lack of modulation may reflect insufficient practice, as de novo learning typically requires longer training bouts than the 90-trial block used here (Telgen et al., 2014; Krakauer et al., 2019; Wilterson & Taylor, 2021; Gastrock et al., 2024). Future work should examine whether extended mirror-reversal training leads to reliable frequency-domain changes. Overall, beta and alpha synchronization during movement preparation are modulated by the error magnitude of the upcoming movement in motor adaptation tasks.

We also find that post-movement feedback processing evolves differently across perturbation types. In the temporal domain, all perturbations elicited a P3 component, but only the fixed and random rotations showed training-related changes. The P3 is linked to cognitive and attentional processing and typically scales with error magnitude (Palidis et al., 2019; Aziz et al., 2020; Verleger, 2020; Reuter et al., 2022). Thus, P3 amplitude should decrease as performance improves. In the fixed rotation, P3 amplitude decreases from early to late training, consistent with reduced errors and a shift toward implicit recalibration observed during motor adaptation (Bastian, 2008; Haith & Krakauer, 2013; Krakauer et al., 2019). In the random rotation, P3 also decreases during late training but to a smaller degree. This may suggest that participants initially attempt to adapt but later disengage due to unpredictability, yet still experience substantial errors to elicit a P3. In the mirror reversal, P3 amplitude remains stable across training. Given participants’ variable compensation and movement errors throughout training, this likely reflects continued reliance on explicit, cognitively demanding strategies. Interestingly, a few milliseconds after the P3 we observe a sustained positivity following small but not large errors. Prior work has shown that this Sustained Late Positivity (SLP) is negatively associated with savings, a marker of explicit learning (Reuter et al., 2020). That is, increased SLP relates to less explicit learning. Because small perturbations typically engage implicit processes (Werner et al., 2015; Neville & Cressman, 2018; Modchalingam et al., 2019), the presence of SLP after small errors aligns with the idea that this component may reflect implicit learning dynamics. Finally, in the frequency domain, effects emerged only during random rotation training. Medial frontal beta was attenuated early in training and resynchronized later, while lateral central beta increased following large compared to small errors. These patterns once again likely reflect participants’ initial attempts to compensate for the unpredictable perturbation before eventually disengaging. Together, these results show that feedback-related neural dynamics differ across perturbation types, with distinct signatures emerging depending on the task demands (e.g. random rotation) and reflecting the relative contributions of implicit and explicit learning processes.

In conclusion, we show that the brain engages distinct preparatory and feedback-related mechanisms dependent on the nature of the visuomotor perturbation experienced. Fixed rotations evoke clear signatures of implicit sensorimotor recalibration across both temporal and frequency domains, whereas mirror-reversal learning shows limited modulation because of the task’s reliance on effortful, explicit contributions to learning and the need for prolonged training with the task. Random rotations elicit initial responses similar to those in adaptation, but is followed by disengagement as the task becomes evidently unpredictable. Given the exploratory nature of our analyses, future work should not only confirm our findings but also examine how EEG markers during mirror reversal training evolve with extended practice, or whether entirely different markers better capture de novo learning. Regardless, our results demonstrate that distinct neural mechanisms underlie different types of motor learning.

## Acknowledgments

This work was supported by NSERC for D.Y.P.H.; NSERC, OGS, and VISTA for R.Q.G. The funders had no role in study design, data collection and analysis, decision to publish, or preparation of the manuscript. We would like to thank Dr. Edward Ody for his assistance with EEG data collection and insights into ERP analyses.

